# Immune genes, *IL1β* and *Casp9,* show sexual dimorphic methylation patterns in the zebrafish gonads

**DOI:** 10.1101/753301

**Authors:** M. Caballero-Huertas, J. Moraleda-Prados, S. Joly, L. Ribas

**Affiliations:** Institute of Marine Sciences, Spanish National Research Council (ICM-CSIC) Passeig Marítim de la Barceloneta, 37–49. 08003. Barcelona

**Keywords:** sexual dimorphism, epimarkers, methylation, gene expression, immune, gonads

## Abstract

There is a crosstalk between the immune and the reproductive systems in which sexual dimorphism is a common pattern in vertebrates. In the last years, epigenetics has emerged as a way to study the molecular mechanisms involved during gonadal development, which are responsible to integrate environmental information that contributes to assign a specific sexual phenotype (either an ovary or a testis). In the fish gonads, it is known of the existence of the reproduction-immune system interactions although the epigenetic mechanisms involved are far to be elucidated. Here, we used the zebrafish (*Danio rerio*) as a model to study the DNA methylation patterns of two well-known innate immune genes: *IL1β* and *Casp9*. DNA methylation levels were studied by a candidate gene approach at single nucleotide resolution and further, gene expression analysis were carried out. Results showed that there was clear sexual dimorphism in the DNA methylation levels of the two immune studied genes, being significantly higher in the testes when compared to the ovaries. In summary, and although much research is needed, here we present two potential candidates as epimarkers with forthcoming applications in the livestock and fish farming production, for example, in immune fish diseases or sexual control programs.

## 1. Introduction

Epigenetic definition has been subjected to controversy (Deans and Maggert, 2015) but, in general terms, it could be defined as the set of reactions conditioned by the cellular environment that are responsible to change DNA activity and gene expression without altering the nucleotide sequence (Moore et al., 2013). These mechanisms enclose changes in histone conformation (e.g., chromatin packaging), DNA methylation and regulation by non-coding RNAs (Qiu, 2006). DNA methylation is the best characterized epigenetic modification functioning on the pre-transcriptional control in several biological processes, such as cell differentiation or genomic imprinting (Jones and Takai, 2001). Epigenetic changes (or epimutations) can be inherited cell to cell through mitotic divisions or throughout subsequent generations by meiotic divisions (Dupont et al., 2009; Hanson and Skinner, 2016). Inheritable epimutations need to occur in the primordial germ cells (PGCs) during embryo formation, which in humans appeared during 6-18 weeks of gestation (Jirtle and Skinner, 2007) and in zebrafish (*Danio rerio*) approximately around 1,000 cells (Koprunner et al., 2001). In addition, gametogenesis in the gonads, i.e., sperm and oocyte formation, is a sensitive window to promote transgenerational epimutation changes that will be transferred to the following phenotypes (Daxinger and Whitelaw, 2012). Thus, understanding the underlied molecular mechanisms of these epimutations in the gonads needs to be at the forefront of the research.

In the fish gonads, the first study showing the interaction between epigenetics and sexual development was published eight years ago in the European sea bass, *Dicentrarchus labrax*, showing that methylation of the gonadal aromatase (*cyp19a1a*) promoter was altered in the ovaries of adult females previously exposed to high temperature during early stages of development (Navarro-Martin et al., 2011). Since then, other studies, including in this fish species (Anastasiadi et al., 2017; Anastasiadi et al., 2018b) have been have evidenced the important role of the epigenetic mechanisms in the gonads. This is the case, for example, in half-smooth tongue sole, *Cynoglossus semilaevis* (Shao et al., 2014), Nile tilapia, *Oreochromis niloticus,* (Wang et al., 2017) or barramundi, *Lates calcarifer* (Domingos et al., 2018), among others (reviewed in Piferrer et al. (2019), all of them revealing the importance of the methylation processes in reproduction-related genes in the ovaries or in the testes and so pointing out the sexual dimorphic epigenetic patterns.

Sexual dimorphism refers to phenotypic differences between the two sexes within a species. In fish, sexual dimorphism is present in a large number of species being females much larger than males, for example in turbot, *Scophtalmus maximus* (Imsland et al., 1997) or E. sea bass (Saillant et al., 2001) or conversely, males much larger than females, for example in Nile tilapia (Toguyeni et al., 1997) or rainbow trout, *Oncorhynchus mykiss* (Bonnet et al., 1999). Sexual dimorphism not only manifests by morphological traits such as growth, but also by behavior (Magurran and Garcia, 2000) or by molecular mechanisms (e.g., gene expression). Sex-biased gene expression has implications beyond evolutionary biology due to the existence of a predominant expression in one sex (Ellegren and Parsch, 2007). Thus, in human, for example, transcriptomic sex-related differences are more remarkable in the gonads, although other tissues such as brain, muscle or liver also exist (Mele et al., 2015). In fish, transcriptomic studies identifying both the number of expressed genes between ovaries and testes and the interactions of sex-specific pathways have been carried out in several fish species such as zebrafish (Small et al., 2009), turbot (Ribas et al., 2016), guppy, *Poecilia reticulata* (Sharma et al., 2014) or E. sea bass (Ribas et al., 2019).

With clinical purposes, in human, many epigenetic studies address sexual dimorphism, for example, in brain, showing differences in the methylation levels (Matsuda et al., 2012) but the vast majority are focusing on the sex-related differences in diseases responses. It is well documented that woman are more resilient against infectious and some non-infectious diseases such as cancer but at the same time, they are more susceptible to autoimmune diseases (Kirsch-Volders et al., 2010; Ghosh and Klein, 2017). Such research is an emerging field in which the underlying mechanisms of these sexual dimorphic responses are far to be elucidated, though it is known, that the epigenetics mechanisms play important roles (reviewed in Klein and Flanagan (2016).

Gonads are immune-privileged sites as they need to prevent immune responses against meiotic germ cells (Maddocks and Setchell, 1990). Immune cells present sexual steroid receptors not only in mammals (Kovats et al., 2010) but also in fish (Chaves-Pozo et al., 2018). Thus, sexual steroids regulate the immune system in addition to classical reproduction-related functions in vertebrates. In fish, immunotoxicology field has developed more of the research concerning the interactions between the reproduction and the immune systems because large number of endocrine disruptors can mimic sex steroids and in turn, activate the immune system (Milla et al., 2011). Further, expression of immune-related genes during gonadal development have been described in some fish species like gilthead seabream, *Sparus aurata* L. (Chaves-Pozo et al., 2008; Chaves-Pozo et al., 2009) or turbot (Ribas et al., 2016). In sea bream, a protandrous hermaphroditic teleost, a massive infiltration of an immune cell type (acidophilic granulocyte) is required to degenerate the testis for phenotype sex-change (Liarte et al., 2007). In zebrafish, gonads are first developed as an ovarian-like organ to latter half of the population activates apoptotic and tp53 signaling pathways that drive to the testis phenotype (Rodríguez-Marí et al., 2010; Liew and Orban, 2014).

Taken together, there is evidence of immune-reproduction interactions in the fish gonads that are able to maintain physiological and molecular changes. Nevertheless, the role of epigenetic mechanisms in regulating these changes by a sex-dependent manner is unknown. We address this gap by studying the methylation levels in the zebrafish ovaries and testes of two well-known innate immune genes: interleukin 1 beta, *IL1β*, a proinflammatory multifunctional cytokine involved in a variety of cellular activities (Secombes et al., 2011) and caspase 9, *Casp9*, a cysteine proteases that plays an essential role in the mitochondria-mediated cell death pathway (Sakamaki and Satou, 2009). We performed a candidate gene approach method (Methylation Bisulfite Sequencing, MBS) which allowed to analyze at single nucleotide resolution, high coverage and single CpG sites in the desired regions (Anastasiadi et al., 2018b). In parallel, gene expression of these two immune genes were studied together with methylation levels by correlation analysis.

## 2. Material and methods

### 2.1. Fish husbandry

AB zebrafish were housed in the fish facilities at the Institute of Marine Sciences (ICM- CSIC, Barcelona) following standard conditions (Ribas and Piferrer, 2014) with an appropriate density levels (8 fish/L) to avoid induced masculinization (Ribas et al., 2017). Fish were reared in a close-circuit system (Aquaneering, San Diego, CA, USA) in a monitored chamber: photoperiod of 12:12 hours, environmental temperature of 26 ± 1°C and humidity of 50 ± 3%. Physicochemical parameters of the water were monitored daily: temperature (28 ± 0.2°C), pH (7.2 ± 0.5), conductivity (750-900 μS) and dissolved oxygen (6.5-7.0 mg l □^1^). Ammonia, nitrate and nitrite water levels were checked weekly by a commercial colorimetric test (Test kit Individual Aquarium Water Test Kit, API, United Kingdom).

### 2.2 Ethics statement

The animal procedure for the present study was licensed by Bioethical Committee of the Generalitat de Catalunya with the reference code 9977 and approved by the Spanish National Research Council (CSIC) Ethics Committee within the project AGL2013–41047–R. European regulations of animal welfare (ETS N8 123, 01/01/91 and 2010/63/EU) were respected regarding fish rearing and maintenance. Fish facilities in the ICM were validated for animal experimentation by the Ministry of Agriculture and Fisheries (certificate number 08039–46–A) in accordance with the Spanish law (R.D. 223 of March 1988).

### 2.3 Fish sampling

A total of ten females and ten males from adult zebrafish at 90 dpf were euthanized on iced water followed by decapitation, and total weight (precision ± 0.05 g), standard length (SL, precision ± 0.01 cm) were recorded. Under magnifying glass, gonads were sexed, dissected and flash frozen in liquid nitrogen and kept at −80°C for further analysis.

### 2.4 Methylation analysis

Methylation levels of the ovaries and testes were analyzed following the protocol described in (Anastasiadi et al., 2018b). Briefly, genomic DNA was isolated from 20 gonads and a unique library was prepared by pooling the *IL1β* and *Casp9* amplicons for all the samples. For each fish, DNA was extracted from half of the gonad which was lysed by by one microgram (1 μ g) of proteinase K (Sigma-Aldrich, St. Louis, Missouri) overnight at 65°C and the following day, a standard phenol-chloroform-isoamyl alcohol protocol with ribonuclease A (PureLink RNase A, Life Technologies) was performed. Five hundred nanograms of DNA per sample were bisulfite-converted using the EZ DNA Methylation-Direct™ Kit (ZymoResearch; D5023).

### 2.5 Primer design

Primers were designed for bisulfite converted DNA using MethPrimer tool (Li and Dahiya, 2002) with an amplicon size of 459 and 441 for *IL1β* and *Casp9*, respectively. Amplicon regions included the promoter, the first intron and the first exon as much as possible (Figure 1, Table S1). The adaptor sequences for 16S metagenomic library preparation (Illumina) were added to the 5□ ends of the primers designed: Forward-TCGTCGGCAGCGTCAGATGTGTATAAGAGACAG and Reverse-GTCTCGTGGGCTCGGAGATGTGTATAAGAGACAG. The target regions included a total of nine for *IL1β* and 19 for *Casp9* CpGs, respectively. Primer specificities were previously validated by Sanger sequencing of amplicons from a pool of samples.

**Figure 1.**
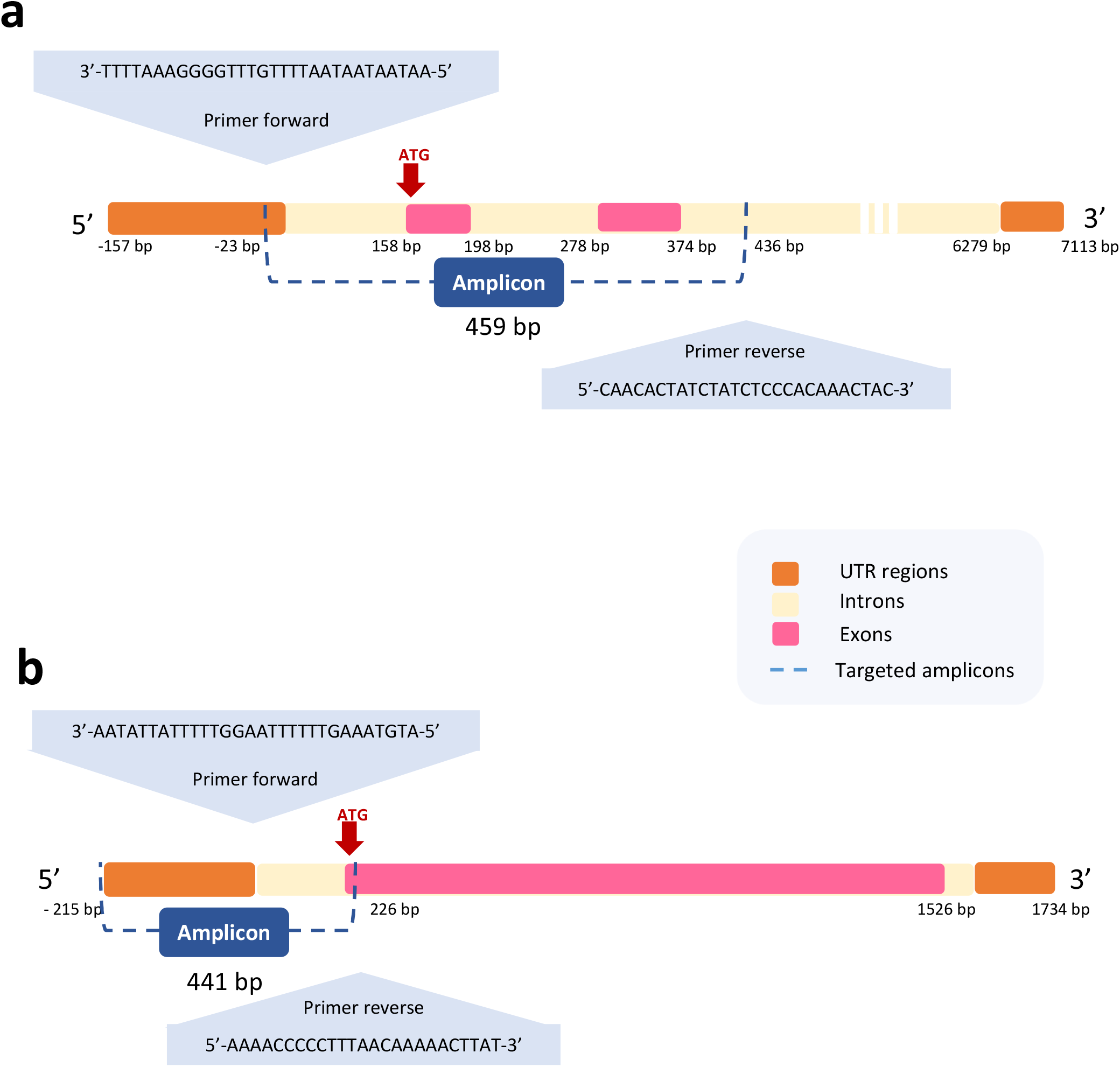
Schematic view of the amplicon region studied by Methylation Bisulfite Sequencing (MBS) method for *IL1β* (A) and *Casp9* (B) genes. The amplicon length for the *IL1β* was 459 bp and for Casp9 441 bp. A total of 9 CpG were studied for *IL1β* and 19 CpG for Casp9. Accession numbers were NM212844.2 and NM001007404.2 for *IL1β* and Casp9, respectively.

### 2.6 Amplification of target regions and library preparation

Targeted-regions were obtained from bisulfited DNA (25 ng per sample) by 30 cycles of PCR using specific primers: 7 min at 95°C, followed by 40 cycles of 95°C for 1 min, annealing temperature (55°C and 54°C for *IL1β* and *Casp9*, respectively) for 2 min and 65°C for 2 min, with a final step at 65°C for 10 min. Obtained targeted amplicons were size-selected and normalized by magnetic beads was (Fisher 09981123) following the protocol described in (Anastasiadi et al., 2018b). PCR products were indexed by Nextera XT index Kit Set A and Set D (Illumina; FC-131– 2001 and FC-131–2004) according to Illumina’s protocol for 16S metagenomic library preparation. PCR products were pooled in an equimolar manner to obtain a single multiplexed library which was sequenced on a MiSeq (Illumina, San Diego, California) using the paired-end 250 bp protocol at the National Center of Genomic Analysis (CNAG, Barcelona).

### 2.7 Biocomputational analysis

Raw data from the sequencer were demultiplexed and Nextera adapters were removed using Trim Galore! software (v. 0.4.5) (Babraham Bioinformatics) and low quality bases were filtered (Phred score < 20). Pre- and post-trim quality control was carried out with MultiQC to ensure that the adapters were cut off correctly (Ewels et al., 2016). Bismark (v.20.0) was used to generate an *in silico* bisulfite converted zebrafish genome (danRer11) as a reference template for alignments (Krueger and Andrews, 2011). Bisulfite conversion efficiency was calculated for each sample with a minimum threshold of 99.0%. One ovarian sample did not passed the minimum thresholds and therefore was deleted. The coordinate positions of all CpG sites were obtained by using “BSgenome.Drerio.UCSC.danRer10” package from BSgenome (Pagès, 2018). The coordinates of the CpG sites of the targeted genes was assessed by BED tools methodology (Quinlan and Hall, 2010) through bedr package (v. 1.0.4) (Haider et al., 2017). From the targeted CpG sites, only those that showed coverage >10 times were retained for statistical analysis. Raw sequencing data was submitted in the NCBI with the accession number: GSE134291.

### 2.8 Gene expression analysis

Gene expression analyses were carried out with half of the gonadal tissue from the same individuals that were used for methylation analysis except for two fish per sex in which no more samples were available. RNA was extracted with chloroform by TRIzol (T9424, Sigma-Aldrich, St. Louis, Missouri) according to manufacturer’s procedures. Quality of the RNA was measured by ND-1000 spectrophotometer (NanoDrop Technologies) and quality was checked on a 1% agarose gel. By following supplier indications, 150 ng of RNA were treated with DNAse I, Amplification Grade (Thermo Fisher Scientific Inc., USA) and retrotranscribed to cDNA with SuperScript III RNase Transcriptase (Invitrogen, Spain) and Random hexamer (Invitrogen, Spain). Quantitative PCR (qPCR) was carried out in triplicate for each sample and two housekeeping genes were used. cDNA samples were mixed with Eva Green chemistry (87H24-001; Cultek), being the cycling parameters: 50°C for 2 min, 95°C for 10 min, followed by 40 cycles of 95°C for 15 s and 60°C for 1 min. qPCRs were run in triplicate in optically clear 384-well plates (CFX-386, Touch BioRad). Dissociation step, primers efficiency curves and PCR product sequencing by Sanger, confirmed the specificity for each primer pair. Primer sequences used for gene expression study are shown in Table S2.

### 2.9 Statistical analysis

All statistical analyses were carried out using R software (v. 3.4.4). Data were expressed as mean ± SEM and the differences were considered significant when *P* < 0.05. Graphs were generated either by using the ggplot2 package (v. 3.1.0) (Wickham, 2016) or Sigma Plot (v. S13.0).

Methylation levels were calculated by averaging the values of each CpG per gene and per sample, whereas for the CpG-specific approach the means per individual CpGs per sample were considered. DNA methylation were evaluated by one-way ANOVA followed by a Tukey’s HSD test. Normality was tested by the Shapiro-Wilk test and, when normality was not followed, methylation proportions logit transformed and a robust non parametric two-way ANOVA was applied by using the WRS2 package (v. 0.9.2) (Mair and Wilcox, 2016).

Statistics on gene expression data was calculated by the 2ΔΔCt method (Livak and Schmittgen, 2001; Schmittgen and Livak, 2008). Data from qPCR were analyzed for normality and homogeneity of variances by using the Kolmogorov–Smirnov and Levene’s test, respectively and significance between two groups was analyzed by t-Student test. For DNA methylation and gene expression correlation analyses, a Spearman’s rank correlation (ρ) test was applied using the corrplot package (v. 0.84) (Wei et al., 2017).

## 3. Results

### 3.1 Sequencing results

The total number of read-pairs that passed the Illumina filter for all 20 samples ranged between 30,000–243,000 read-pairs with an average of 97894.7 ± 47209.8 (mean ±SD) and the bisulfite conversion rate was >99.5% for all samples producing 0.936 Gb of data. MultiQC summary confirmed that sequence quality post trimming passed the quality controls and were within the accepted thresholds.

### 3.2 Methylation patterns show sexual dimorphism

Global methylation were significantly higher in testes than ovaries of adult zebrafish for *IL1β* (*P* < 0.001) and *Casp9* (*P* < 0.05) genes (Fig. 2A and Fig.3A). The percent levels for *IL1β* were 89.7 ± 3.8 and 61.4 ± 5.5 in testes and ovaries, respectively and for *Casp9* were 0.62 ± 0.04 and 0.56 ± 0.12 in testes and ovaries, respectively.

**Figure 2.**
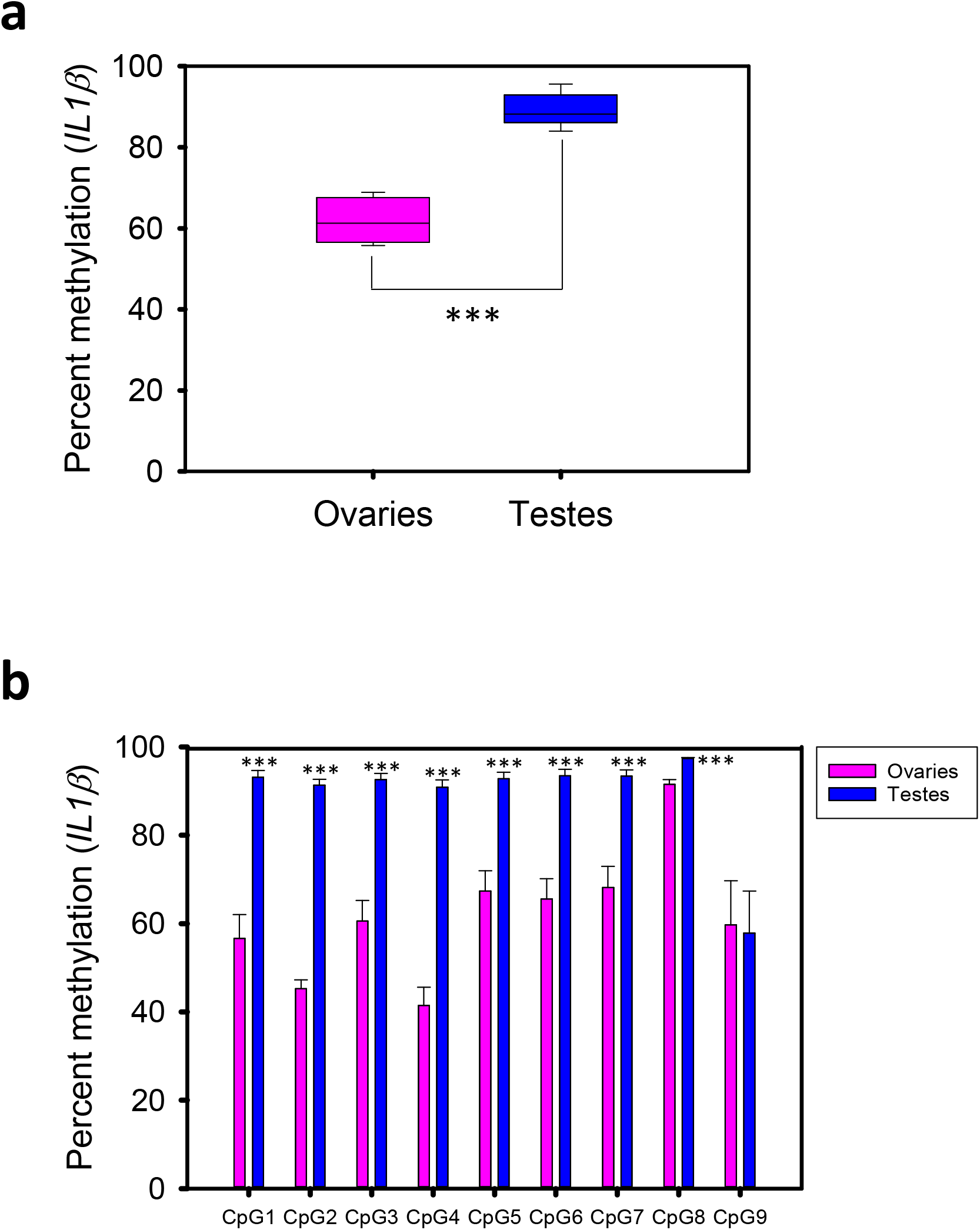
Methylation levels of *IL1β* gene in adult zebrafish gonads. (a) Global DNA methylation percent found in ovaries and testes. (b) DNA methylation percent for each of the nine CpG analyzed. Number of individuals studied; N = 9 ovaries and N = 10 testes. Data shown as mean ± sem. Significant differences *** (P < 0.001).

**Figure 3.**
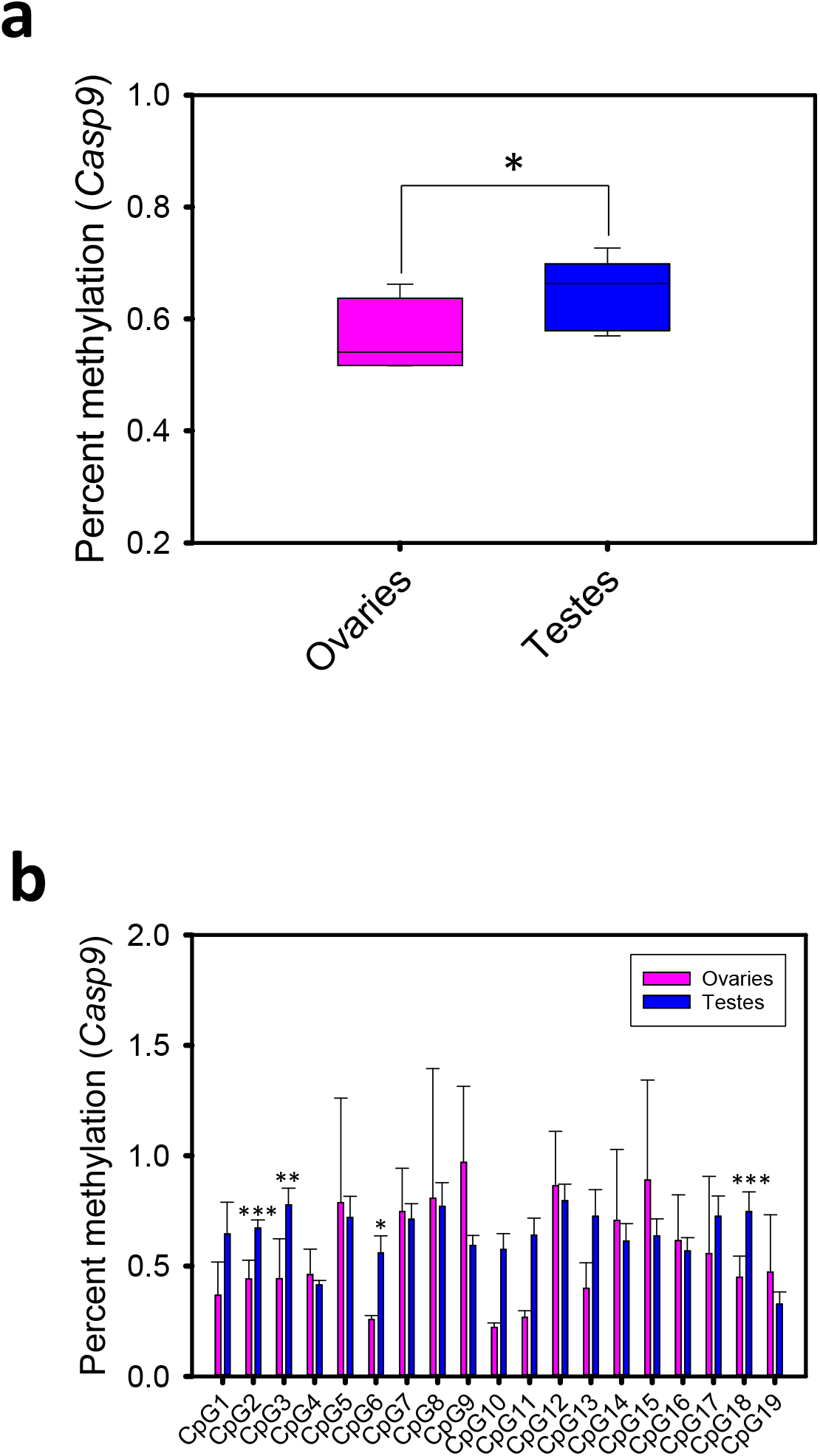
Methylation levels of *Casp9* gene in adult zebrafish gonads. (a) Global DNA methylation percent found in ovaries and testes. (b) DNA methylation percent for each of the 19 CpG analyzed. Number of individuals studied; N = 9 ovaries and N = 10 testes. Data shown as mean ± SEM. Significant differences * P<0.05, ** P < 0.01, ***P < 0.001.

In *IL1β*, eight out of nine analyzed CpGs showed significant differences of the methylation levels between ovaries and testes (*P* < 0.001) (Fig.2B). Only CpG9 was not significant. In *Casp9*, four out of nineteen CpGs showed significant difference in the methylation levels between ovaries and testes which significant levels varied among CpG positions: CpG2 and CpG18 *P* < 0.001; CpG3 *P* < 0.01 and CpG 6 *P* < 0.05 (Fig.3B).

### 3.3 Gene expression

The fold change of *IL1β* expression in the testis was seven times higher than that of ovary (*P* < 0.01) in adult zebrafish (Fig.4A). However, no significant differences in gene expression were found for *Casp9* gene when testes and ovaries were compared (Fig.4B).

**Figure 4.**
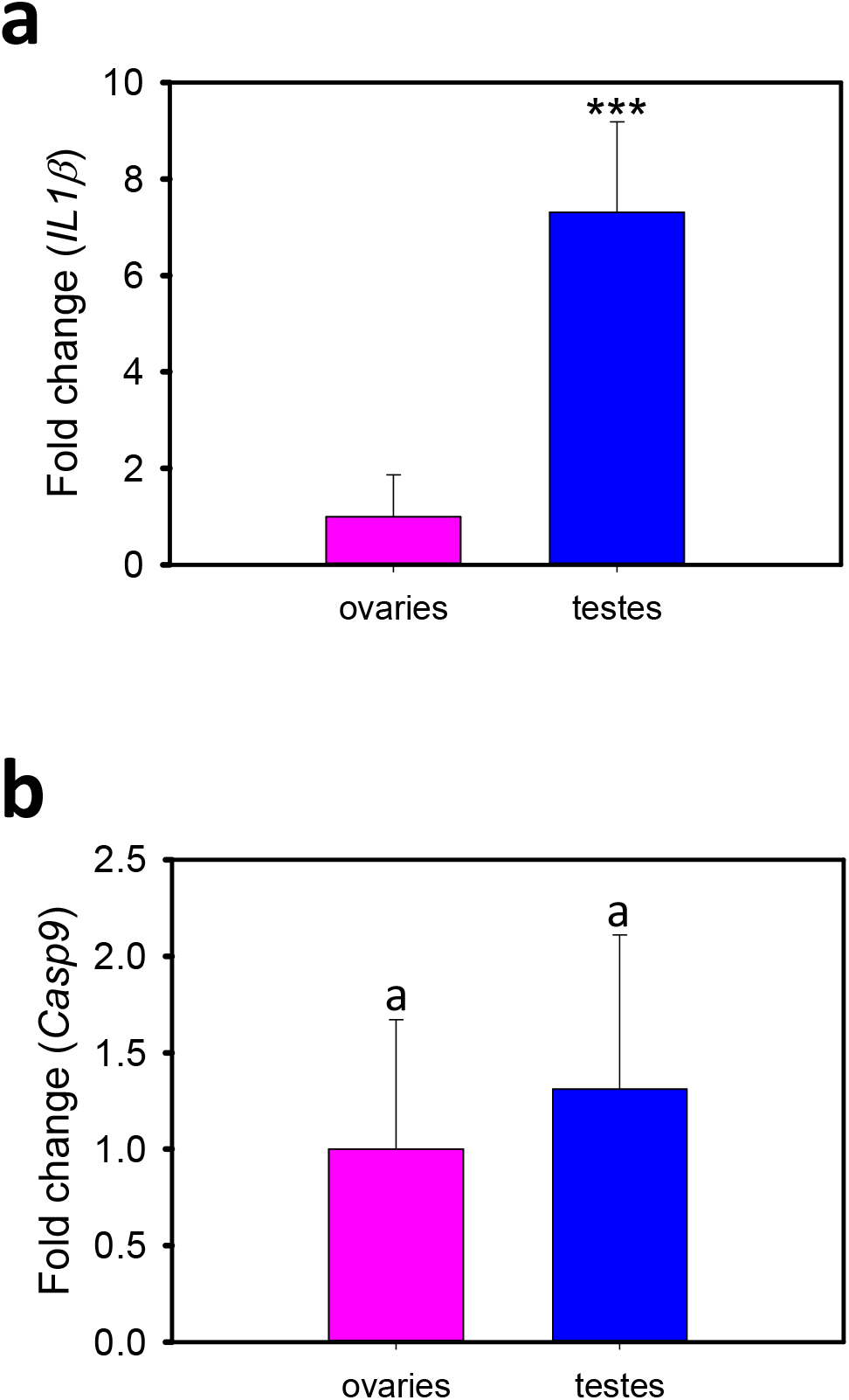
*IL1β* and *Casp9* expression profiles in adult zebrafish gonads. Data shown as mean ± SEM of fold change using control values set at 1. Sample size *n* = 10 gonad per sex. Significant differences between sexes were analysed by Student’s *t* test, *** *P* < 0.001.

### 3.4 Correlations methylation vs. gene expression

Negative correlation patterns between methylation and gene expression levels were observed for both *IL1β* and *Casp9* genes in testes and ovaries of adult zebrafish, though no significance differences were obtained (Fig. 5). The strongest correlation was found in the testes for *IL1* gene (ρ = −0.5).

**Figure 5.**
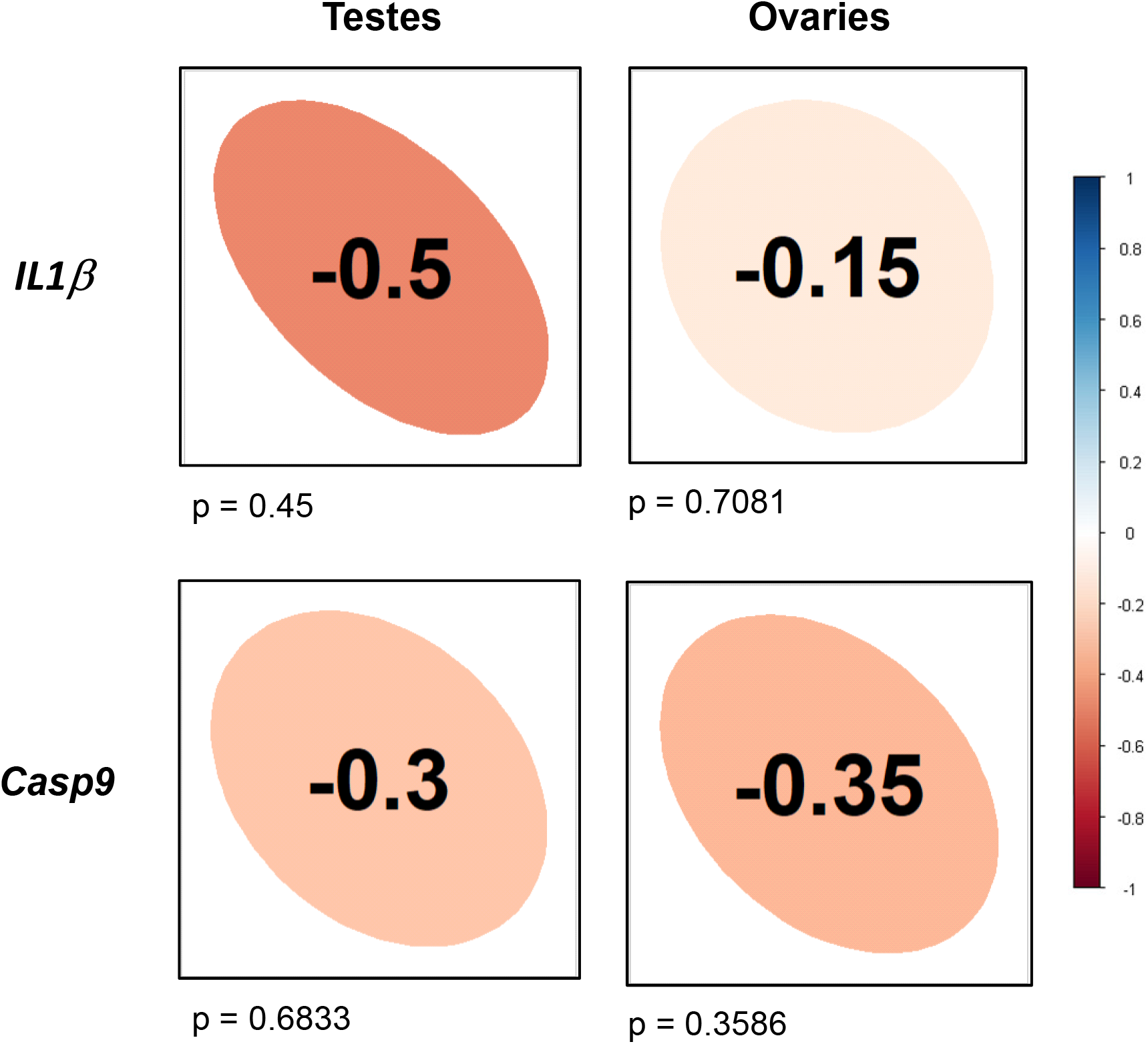
Correlations DNA methylation and gene expression of *IL1β* and *Casp9* genes. Correlations are shown by Spearman’s rank correlation coefficient (ρ). The direction of the long axis of the ellipses and the color indicate the type of correlation: negative is shown in red and positive is in blue shade. The short axis of the ellipse and the intensity of the color are proportional to the correlation coefficients. Significant differences are considered when *= *P* < 0.05; **= *P* < 0.01; *** = *P* < 0.001.

## 4. Discussion

The main immune signaling pathways are conserved in fish (Stein et al., 2007) in which the innate response is the essential immunity to cope immune challenges (Press and Evensen, 1999) although evidence of the required role of the adaptive immunity have been emerged in the last years (Secombes and Belmonte, 2016). The innate response is the earliest immune mechanism characterized by being nonspecific and therefore not depending upon previous recognition of the surface structures of the invader (Tort et al., 2003). It has also the advantage of being inducible by external molecules but at the same time is constitutive and reacts within a very short time scale thorough inflammatory pathways (Bayne and Gerwick, 2001). Interleukins constitute a large group of cytokines involved in the specific immune defense by recruiting leukocytes to a site of infection to initiate inflammation (Brocker et al., 2010). *IL1β*, together with IL6 and tumor necrosis factor alpha (TNFα), are one of the main actors involved in the first inflammatory response (Smith et al., 2000). *IL1β*, once it is cleaved by caspase 1 into a mature form, it is able to bind to its receptor leading the activation of the nuclear factor (NF)-kβ pathway (Martin and Falk, 1997). Caspases, including *Casp9*, are a family of protease enzymes that play essential roles in the apoptotic pathways and in the inflammasome of the cells (McIlwain et al., 2013; Shalini et al., 2015).

In vertebrates, and in some invertebrates such as parasites, reproductive and immune systems are interconnected playing sexual-dependent molecular functions (Folstad and Karter, 1992; Nunn et al., 2009). In human, testes contain immune cells such leukocytes and macrophages in the interstitial compartment that are involved in gametogenesis and Leyding cell development (Li et al., 2012). *IL1α* is mostly produced by spermatocytes but also by Sertoli cells and its production is stage-specific (Haugen et al., 1994; Lie et al., 2012). In human ovaries, *IL1α* mediates inflammatory processes during ovulation, fertilization and oocyte implantation (Polan et al., 1991; Garcia-Velasco and Arici, 1999) and it is highly expressed in ovarian cancers (Aggarwal and Gehlot, 2009). In sea bream gonads, *IL1β* was found in the cytoplasm of spermatogonia, spermatocytes and oogonia and its expression was higher during reproductive cycles suggesting a role in promoting proliferation in gametogenesis and spermatocyte differentiation (Chaves-Pozo et al., 2008). The two cytokines, *IL1β* and *TNFα* modulated the *in vitro* testosterone production in goldfish testis indicating their role in regulating sex steroids (Lister and van der Kraak, 2002). In the present work, we report a higher expression of *IL1β* in testes when compared to ovaries in adult zebrafish according to what previously found by Lee et al. (2017), thus indicating that *IL1β* is expressed in a sex-dependent manner in fish. We did not find significant difference in *Casp9* expression between testes and ovaries although higher expression in testes was also observed. In contrast, in protogynous black sea bass (*Centropristis striata*) ovaries expressed *Casp9* twice as much as testes, although gene expression patterns related to atresia differs among gonochoristic and hermaphroditic species (Breton et al., 2019). It is described that in zebrafish, during gonad development, activation of apoptotic pathways are required for testes development (Liew and Orban, 2014), including specifically the expression of *Casp9* (Pradhan and Olsson, 2014), thus we might have observed the remaining activation of the pro-male machinery during testicular development.

Epigenetic modifications occurred in the host genome by pathogens after infections altering the transcription and the corresponding signaling pathways of the host (Marr et al., 2014). The understanding of epigenetic host-pathogen interactions is a flourishing research field to untangle the sophisticated cellular strategies (Gomez-Diaz et al., 2012). In humans, plenty of studies give evidence of rapid change in DNA methylation of the innate immune cells to face infections (Sinclair et al., 2015; Wiencke et al., 2016; Pacis et al., 2019) and the importance of the epigenetic mechanisms in reprograming the immunity (Netea et al., 2016). In contrast, less data is found in fish. Recent findings in guppy (*Poecilia reticulate*) showed changes in DNA methylation dynamics after host–parasite interactions (Hu et al., 2018) and in Atlantic salmon (*Salmon salar*) showed that stress response together with immune challenges, altered transcriptome and methylome of the gills through immune responses (Uren Webster et al., 2018). In zebrafish, viral infections showed that histone modification was able to increase methylation levels of the gene promoters associated with innate immune response in head kidney, liver, spleen and heart (Medina-Gali et al., 2018). Surprisingly, to our knowledge, no data on epigenetic mediating infections in the immune system in the fish gonads are currently found.

It is estimated that in zebrafish, global methylation in sperm is nearly 95% of CG dinucleotides are methylated while in oocytes are ∼75% methylated (Jiang et al., 2013; Potok et al., 2013). Therefore, it may be said that the product of mature germ cells already possess methylation sexual dimorphism. Here, by a candidate gene approach method, we found higher methylation levels in the two studied immune genes (*IL1β* and *Casp9*) in the testes when compared to ovaries thus evidencing sex-specificity of the methylation programs in the fish gonads. Likewise, significant differences were found in specific CpG sites for each of the targeted regions, in which eight out of nine and four out of nineteen of the studied CpGs for *IL1β* and *Casp9*, respectively, showed significant differences in the methylation levels between the two sexes. Global DNA methylation levels between these two immune cells differ in magnitude being ∼73% and ∼0.62% (percent methylation mean in ovaries and testes) in *IL1β* and *Casp9*, respectively. Although low levels were found in *Casp9* gene, percent difference between ovaries and testes of the four CpG significantly methylated was ∼2%. In a similar manner, differences in the methylation levels (5.2%) were enough to observed epigenetic transgenerational effects in the insulin growth factor (IGF) 2 promoter in the siblings of those mothers famished in the Second World War (methylation levels for IGF2 ranged from 0.517-0.488 in all studied groups) (Heijmans et al., 2008). Thus, low but significant changes in DNA methylation levels might induce epigenetic changes.

Analysis of gene expression and methylation levels showed negative, although not significant, correlations among the four studied comparisons. Conventionally, higher methylation levels in the gene promoters have been associated with transcriptional silencing (Illingworth and Bird, 2009; Straussman et al., 2009). However, gene expression inhibition have been related to other genomic features like the first exon and the first intron of the gene body not only in human (Blattler et al., 2014) but also in fish (Anastasiadi et al., 2018a). Further, the approach here performed for studying the methylation levels is limited to amplicons about ∼500 bp, that even with the intention of include promotor, first intron and first exon in the targeted regions, the entire gene body and possible enhancers involved in the gene regulation were not fully studied. Thus, to tackle this physical barrier, wide genomic approaches need to be performed.

In tilapia, genome-wide methylation analysis identified sexually dimorphic methylated regions along specific genomic regions (Wan et al., 2016) as well as in zebrafish brain, global methylation differences were found between males and females (Chatterjee et al., 2016). Thus, both studies, pointed out the importance of the sexual dimorphic methylation patterns in fish. Our results supports that sexual dimorphism defined the methylation patterns in two immune genes in the gonads and so the sex-biased differences need to take into account when analyzing the immune responses in fish. In fact, human research on sex-based differences in immune responses are bringing to light the importance of considering sex when analyzing the data in such a way that funding agencies and journals have launched policies to analyze and publish research based on sex and gender in biomedical sciences (Klein and Flanagan, 2016). Thus, sexual dimorphic differences need to be included for good data interpretation.

Over the last few years, epigenetic marks have been implemented to predict, among others, phenotypic features. This is the case, for example, in the identification of clinical biomarkers to predict cancer predisposition or other diseases (Jubb and Harris, 2010; Tabernero et al., 2015). In animal production, epigenetic biomarkers to predict the sex of the offspring in E. sea bass has been stated as an important step towards improving sex control programs in aquaculture farms (Anastasiadi et al., 2018b). The present study shows, for the first time, the presence of sexual dimorphism in the DNA methylation profiles of two immune genes in the fish gonads that might be useful, for example, to develop sexual epimarkers but also to other applications such as fish diseases research. Further studies need to cover the methylation patterns of these two immune genes when fish are subjected to immune challenges and to see whether dimorphic sex-specific patterns are present when facing immunological diseases.

## Supplementary table legends

**Table S1.** Information related to the amplicons studied by Methylation Bisulfite Sequencing (MBS)

**Table S2.** Primer information for gene expression analysis of *IL1β* and *Casp9* genes

